# Allelic variation in the wheat homolog of topoisomerase *II* is associated with crossover rate

**DOI:** 10.1101/2023.07.06.548017

**Authors:** Katherine W. Jordan, Monica Fernandez-de Soto, Abraham Korol, Alina Akhunova, Eduard Akhunov

## Abstract

Meiotic recombination is a fundamental biological process that impacts genetic diversity and response of variants to selection. The increase in crossover (CO) frequency by modulating the activity of meiotic genes has been suggested as potential means to improve the efficiency of selection and mitigate the negative impact of linkage drag in crop improvement programs. In this study, we used the wheat nested association mapping (NAM) population to map QTL associated with CO rate and harboring *TaTOPII-A1*, a gene encoding topoisomerase II that is involved in the regulation of meiosis. An amino acid changing SNP (T/C) located in exon 19 of *TaTOPII-A1* with a predicted functional effect showed an association with CO rate. The T-allele of *TaTOPII-A1* was associated with a statistically significant 5.9 % increase in COs across several NAM families. The involvement of *TaTOPII-A1* in processes affecting CO rate in wheat was confirmed using a *TaTOPII-A1* mutant. A strong-effect missense mutation in the *TaTOPII-A1* coding region resulted in 1.4-fold increase in the genetic map length and 53% increase in CO rate compared to the wild-type allele. These results demonstrate that functional mutations in *TaTOPII-A1* can lead to increased CO frequency. The mutant and natural alleles of *TaTOPII-A1* identified in our study could serve as new sources of recombination-promoting genes to manipulate CO rate in wheat and possibly other crops.

## Introduction

Meiotic crossovers (COs) lead to the exchange of genetic information between parental chromosomes and new combination of allelic variants in progeny. Distribution of COs along chromosomes is skewed towards the ends of chromosomes. The prevalence of COs in the distal chromosomal regions is especially prominent for the large genomes of major crops, including barley and wheat^1,2^. This CO patterning complicates selection of beneficial alleles in crop breeding and reduces the resolution of genetic mapping in these regions. Understanding mechanisms driving CO distribution and frequency will be critical for developing strategies to control meiotic recombination and improving the efficiency of breeding approaches when additional rounds of recombination are required to break linkage blocks.

Meiotic recombination involves the formation of double strand breaks (DSBs), which are repaired as crossovers (CO) or non-crossovers (NCOs), with only limited number of DSBs resulting CO ^3^. The CO events are defined by the concerted action of two pathways, inference sensitive pathway dependent on a group of ZMM proteins (class I COs) and interference insensitive MUS81 pathway (class II COs)^3^. In class I COs, which account for up to 85% of all COs, the occurrence of a COs result in suppression of COs in close proximity, with interreference declining with increase in the distance between COs. The functional screens uncovered a role for various classes of genes in the meiotic recombination process and helped to identify those that negatively impact CO rate ^4–6^. Mutations in these anti-crossover genes (e.g. *RECQ4, FANCM*,) were shown to result in increased CO frequency ^7^. For example, loss-of-function mutations in *RECQ4* or *FANCM* have resulted in 2-fold increase in CO rate in pea and rice ^4^. In barley, a mutation in *HvRECQL4* gene also led to 2-fold increase in COs^8^. A significant but less dramatic 31% increase in CO rate was overserved in the double and triple mutants of *FANCM* in tetraploid and hexaploid wheat, respectively ^9^.

In yeast, topoisomerase II (*TOPII*) regulates the distribution and frequency of COs, with the reduction of *TOP II* activity being associated with an increased CO rate^10^. Contrary to these trends, the patterns of CO distribution and their count in the *top II* mutants of *Arabidopsis* were not significantly different from wild-type genotypes ^11^, suggesting that the role of *TOPII* in CO formation in plants and yeast could differ. However, the genetic mapping of meiotic genes in wheat points to a possible association between *TOPII* and natural variation in CO count ^12^. Using a nested-association mapping population, a total of 27 QTL regions associated with CO number variation were detected in wheat^12^, with one QTL on chromosome 6A overlapping with the *TOPII* gene homolog (henceforth, *TaTOPII-A1*). Here, wheat NAM population data, deep exome re-sequencing of NAM founders and EMS mutagenesis of *TaTOPII-A1* gene were combined to investigate the relationship between variation in the coding sequence of *TaTOPII-A1* and CO rate. Our results show that natural variation in *TaTOPII-A1* correlates with the total number of COs in wheat in multiple genetic backgrounds. Mutagenesis confirms the involvement of *TaTOPII-A1* in defining CO number and suggests that both natural and mutant variants of *TaTOPII-A1* can be used for manipulating CO frequency in wheat.

## Results

### Natural Variation in the *TaTOPII-A1* gene in hexaploid wheat

Previously, 27 unique QTL regions were identified across 28 families comprising the spring wheat NAM population that affect natural variation in CO incidence^12^. A QTL detected on chromosome 6A in the NAM18 family, spanned the region containing the phylogenetically conserved wheat homolog of topoisomerase II (*TaTOPII-A1*; gene model TraesCS6A02G268600). The *TaTOPII-A1* gene has homoeologous copies on chromosomes 6B (TraesCS6B02G295700) and 6D (TraesCS6D02G246300), and 5 paralogs in the A genome on chromosomes 1A, 7A, 5A and 3A. No QTL associated with CO frequency in the NAM population were detected in the syntenic regions of homoeologous chromosomes 6B and 6D^12^. The *TaTOPII-A1* coding region includes natural variation (SNP C/T) at the wheat 90K array marker IWB34510^13^ between the common parental cultivar Berkut and line CI 15144. This missense SNP is located at position 1,241 within exon 19 and causes amino acid change from proline to leucine (P1241L), which is predicted to have a strong effect on protein function based on the SIFT score of 0.05^14^.

A single marker test within the NAM18 family found that progeny carrying the T-allele averaged 2.9 more COs than the progeny carrying the C-allele (Figure 1a, Table 1; p-value = 0.041, *t*-test). This SNP also naturally segregates in three other NAM families, NAM19, NAM20, and NAM24. Grouping RILs from the NAM24 and NAM20 families based on the genotype of the IWB34510 marker showed a similar trend, with the T-allele RILs having 3.4 more COs in each family compared to RILs carrying the C-allele (Table 1; *t*-test p-values: NAM24 = 0.049, NAM20 = 0.027). To gain more power, the family effects were removed and the residuals of the fitted model across all four families were compared. This analysis showed a significant increase in the CO incidence (2.8 COs) in the group of RILs with the T-allele compared to RILs with the C-allele (Figure 1b, Table 1; p-value = 9.8 × 10^−4^, ANOVA). These results suggest that natural variation at *TaTOPII-A1* correlates with the number of COs across multiple genetic backgrounds.

**Figure 1.**
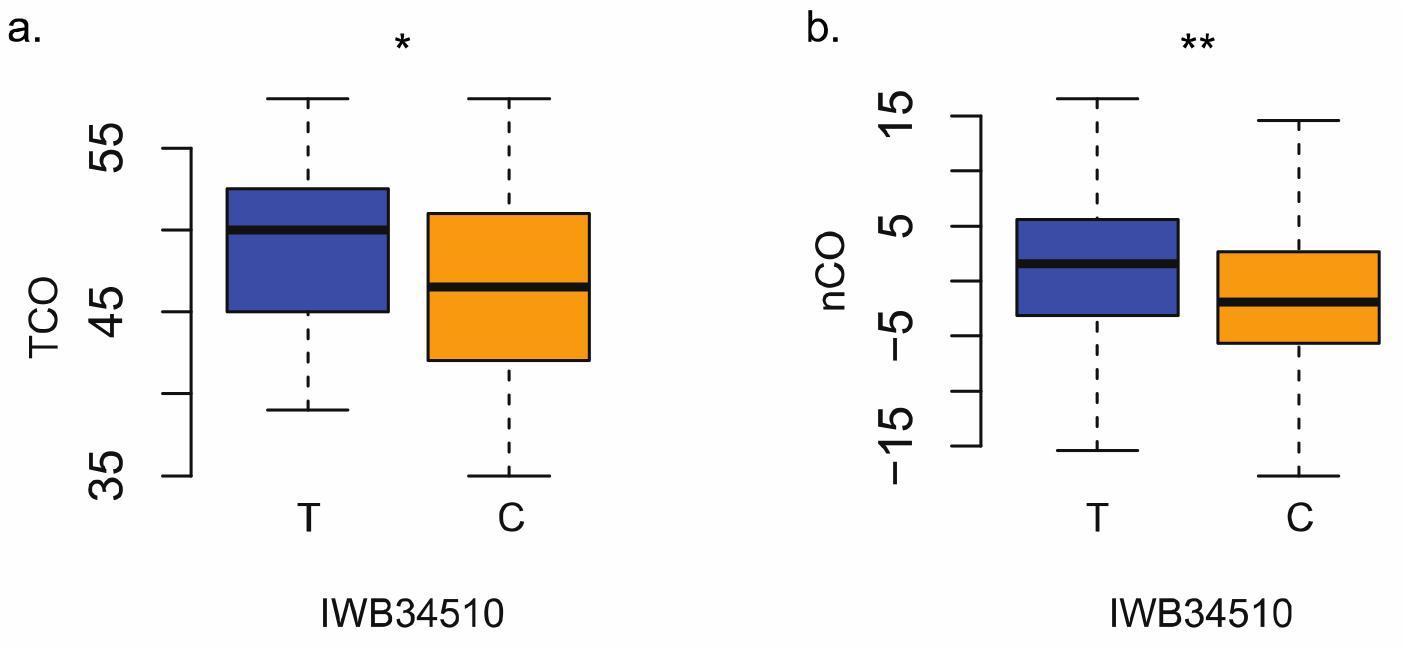
Relationship between the natural variation in the *TaTOPII-A1* gene and COs in the NAM population. **a)** Family NAM18 RILs compared at the IWB34510 marker, lines carrying the T-allele (blue) average 2.9 COs more than lines carrying the C-allele (orange). **b)** Normalized CO (nCO) number for all RILs segregating for the IWB34510 marker, RILs carrying T-allele (blue) average 2.8 CO more than the RILs with C-allele (blue), * *p-value* < 0.05, ** *p-value* < 0.01. Normalized CO (nCO) rates represent the residual value of the individual mean after subtracting the family effect.

**Table 1.**
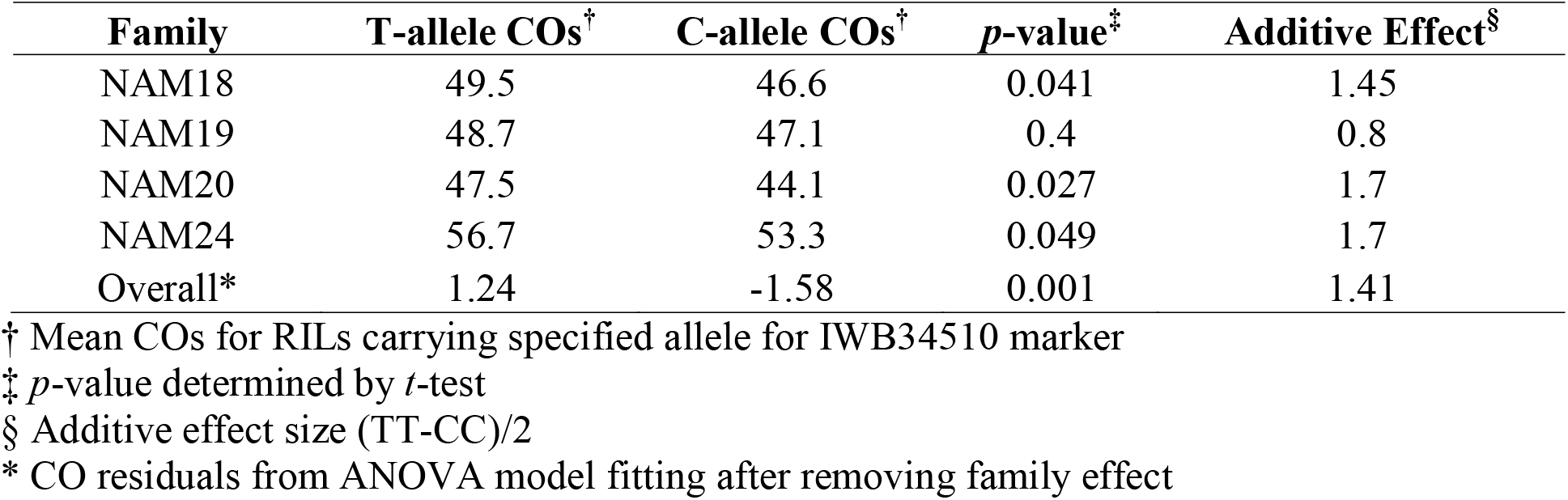
Number of COs in RILs from four NAM families grouped based on the IWB34510 marker genotype.

| Family | T-allele COs<sup>†</sup> | C-allele COs<sup>†</sup> | *p*-value<sup>‡</sup> | Additive Effect<sup>§</sup> |
| --- | --- | --- | --- | --- |
| NAM18 | 49.5 | 46.6 | 0.041 | 1.45 |
| NAM19 | 48.7 | 47.1 | 0.4 | 0.8 |
| NAM20 | 47.5 | 44.1 | 0.027 | 1.7 |
| NAM24 | 56.7 | 53.3 | 0.049 | 1.7 |
| Overall* | 1.24 | -1.58 | 0.001 | 1.41 |

<sup>†</sup> Mean COs for RILs carrying specified allele for IWB34510 marker
<sup>‡</sup> *p*-value determined by *t*-test
<sup>§</sup> Additive effect size (TT-CC)/2
\* CO residuals from ANOVA model fitting after removing family effect

### Epistatic interactions with the natural variants of *TaTOPII-A1* alleles

The lack of a significant association between the IWB34510 marker and CO rate in the NAM19 family suggests that an interaction with the genetic background could modulate the IWB34510 marker effects. We tested for epistatic interactions between IWB34510 and 30 genetic markers associated with variation in CO count in the NAM population^12^. For each segregation region, we selected most significant markers for CO count passing Bonferroni corrected *p*-value threshold of 1.5 × 10^−3^. We detected four markers showing a significant epistatic interaction with IWB34510 (*p-value* ≤ 0.01, ANOVA, Figure 2, Table 2, Supplementary Table1). The two markers that produce the largest epistatic effects on IWB34510 are presence-absence GBS tags that were identified using joint mapping across NAM population^12^. These two markers have opposite effects on the IWB34510 locus. The presence variant at the SW_tag_256152 locus, enhances the effect of the T-allele by more than 3-fold, (Fig. 2; Table 2; *p-value* = 0.0012). The absence variant at the SW_tag_177026 locus suppresses the T-allele effect by more than 2-fold (Table 2, *p-value* = 0.008). In addition to these two PAV variants, two SNP markers from the 90K wheat array^13^ showed an evidence of interaction with the *TaTOPII-A1* locus. The C1 allele of the IWB75092 marker increased the effect of T-allele 2.6 times, where C2 nearly abolished the effect of T-allele on CO count (*p-value* = 0.001). Likewise, in the presence of the C1 allele at marker locus IWB12235, the effect of T-allele was enhanced 1.5 times (*p-value* = 0.01). Of these four interactions, the regions affecting *TaTOPII-A1* locus were located on chromosomes 2B, 3B, and 6A. The magnitude of these interaction effects suggests that genetic background is capable of both enhancing and suppressing the effect of the *TaTOPII-A1* locus on CO rate.

**Figure 2.**
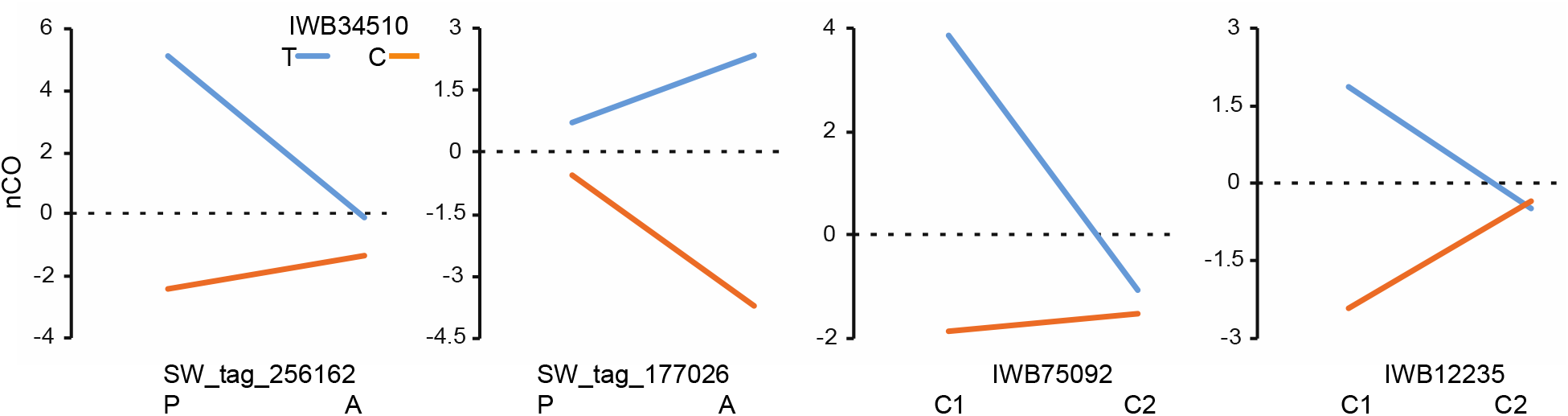
Epistatic interactions with the IWB34510 SNP site. Each panel represents a previously identified QTL region for the total number of COs (TCO)^12^. For each comparison the normalized CO (nCO) is represented by the T-allele (blue) and C-allele (orange) of IWB34510 in the background of the epistatic marker. For the interacting presence/absence variants, presence (P) and absence (A) are indicated on *x*-axis. Normalized CO (nCO) rates represent the residual value of the individual mean after subtracting the family effect.

**Table 2.** Epistatic Interactions with the IWB34510 SNP site.

| Marker Type | Chr: Pos | Phenotypic enhanceme nt† | Phenotypic Suppression‡ | Fold Change§ | Enhancing Allele | p-value |
| --- | --- | --- | --- | --- | --- | --- |
| SpringWheatNAM_tag_256152 (GBS PAV) | 2B: 581 Mb | 7.54 | 1.25 | 2.7 | presence | 0.001 |
| SpringWheatNAM_tag_177026 (GBS PAV) | 3B: 549 Mb | 6.07 | 1.28 | 2.2 | absence | 0.008 |
| IWB75092 (90K SNP) | 6A: 606 Mb | 5.73 | 0.46 | 2.0 | C1 | 0.001 |
| IWB12235 (90K SNP) | 2B: 18.2 Mb | 4.27 | -0.15 | 1.5 | C1 | 0.01 |

† Difference in CO# in enhancing background for T allele ( $CO_{(T,Y1)} - CO_{(C,Y2)}$ ); $Y1 > Y2$
‡ Difference in CO# in suppressing background for T allele ( $CO_{(T,Y2)} - CO_{(C,Y1)}$ ); $Y2 > Y1$
§ Fold change of enhancing effect with respect to T allele original effect size (2.8 CO)

### Effect of an EMS-induced mutation in *TaTOPII-A1* on CO rate in cv. Kronos

To test whether the observed relationship between natural variation in *TaTOPII-A1* and CO rate is simply correlated response or caused by mutation in the *TaTOPII-A1* coding sequence, we utilized a mutant identified in the EMS population generated for tetraploid wheat cv. Kronos^15^. EMS line 2306 carries a heterozygous nonsynonymous mutation A901T with the predicted deleterious effect on protein function (SIFT score = 0.02). Using KASP marker designed for this site, we identified two M_4_ generation mutants of cv. Kronos that were heterozygous at the A901T mutation site. These plants were used to produce two M_5_ families including 56 (family 1) and 51 (family 2) lines (Fig. 3a). The M_5_ plants from both families were genotyped using the same KASP assay to assign lines into the heterozygous and homozygous wild-type or mutant groups. Segregation ratios at the A901T site followed the expected 1:2:1 ratio in both families (*p-value* = 0.62 family 1, *p-value* = 0.16 family2, χ^2^ test). Further, homozygous wild-type and homozygous mutant plants from each family were genotyped using the KASP assays developed for random set of 112 EMS mutations identified by whole exome capture of M_2_ generation plants^15^ (see Methods and Supplementary Table 2). Due to the low density of EMS mutations still segregating in M_4_ generation plants, only a small fraction of KASP markers were useful for counting COs. To increase marker density, the same set of M_5_ generation lines was genotyped using the GBS approach. Though GBS is not ideal method for discovering EMS-induced mutations in the wheat genome, with the density of 2-3 mutations per 100 Kb^16^, it was cost-effective option to improve marker coverage in our populations to better assess CO rates. The genotyping data for informative KASP and GBS markers were merged and analyzed to compare the genetic map lengths for wild-type and mutant groups. Considering the low-density of EMS mutations that could be discovered using these two methods of genotyping, our analyses were focused on the subsets of chromosomes that had at least 5 markers mapped to the reference genome. For family 1, 45 informative genetic markers were selected for chromosomes 2B, 3B, and the short arms of chromosomes 6A and 6B (Supplementary Figure 1). For each chromosomal comparison the map length estimated using data from mutant group was longer than the map length based on wild-type dataset. On average, mutant map sizes were increased 1.7-fold compared to wild-type chromosome sizes (Table 3). Similar results were found in family 2 for chromosomes 2A, 4A, 4B, 5A, and 7B, whose map lengths increased on average by 1.3-fold compared to the wild-type maps (Supplementary Figure 2). No detectable increase in the map length was found for chromosome 5B. Overall, summing genetic map lengths in both families, mutant *TaTOPII-A1* allele increased genetic map distance by 1.4-fold compared to the map length in the wild-type population (Table 3).

**Figure 3.**
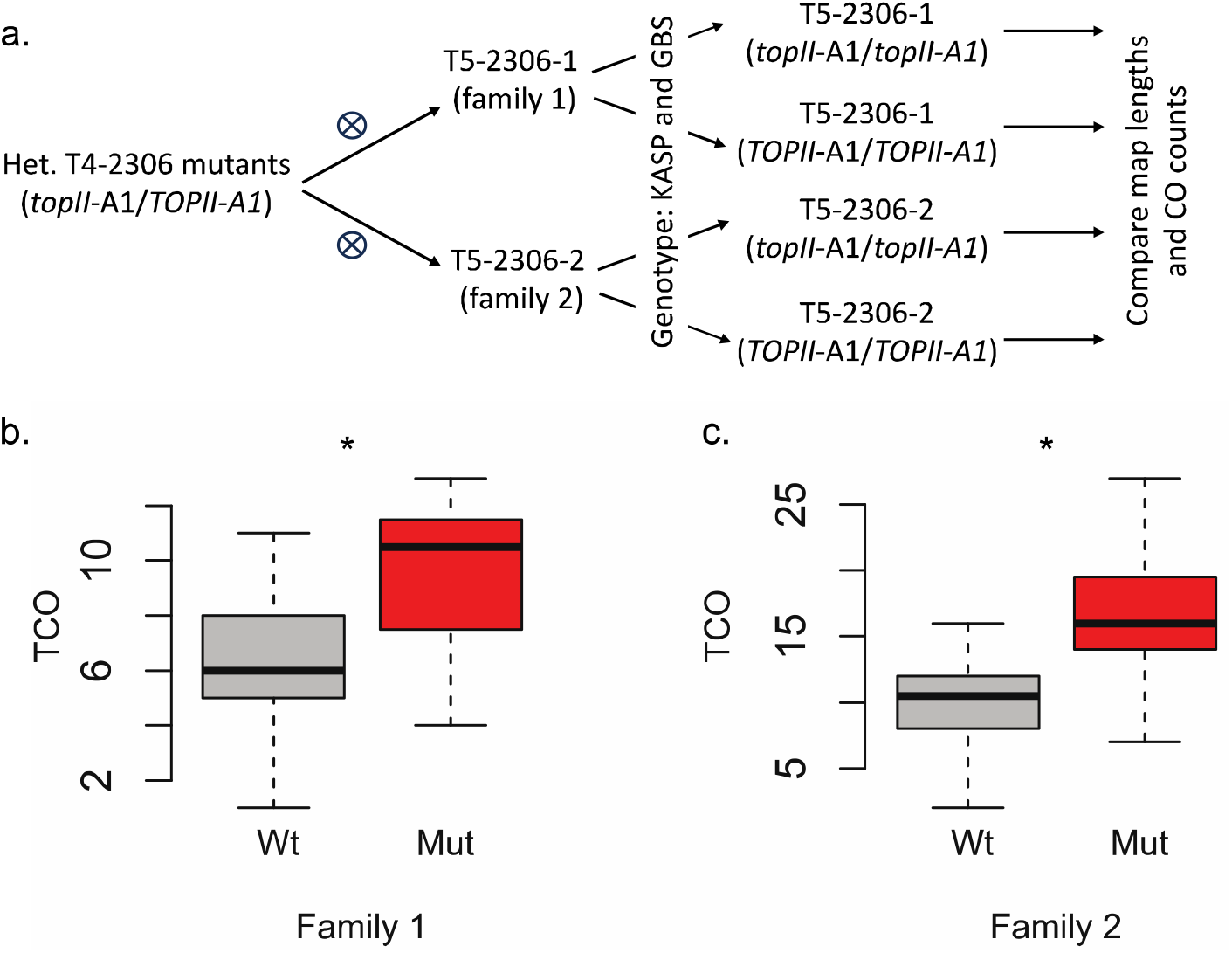
Effects of A901T mutation on CO rate in tetraploid wheat cultivar Kronos. **a)** A scheme showing development of populations for assessing the impact of *TOPII-A1* on COs. **b)** Family 1 comparing homozygous mutant and wildtype progeny, resulted in average increase of 3.3 CO in mutants across 3.5 chromosomes, * *p-value* ≤ 0.02. **c)** Family 2 resulted in average increase of 5.7 CO in mutants across 5.5 chromosomes, * *p-value* ≤ 0.02.

**Table 3.**
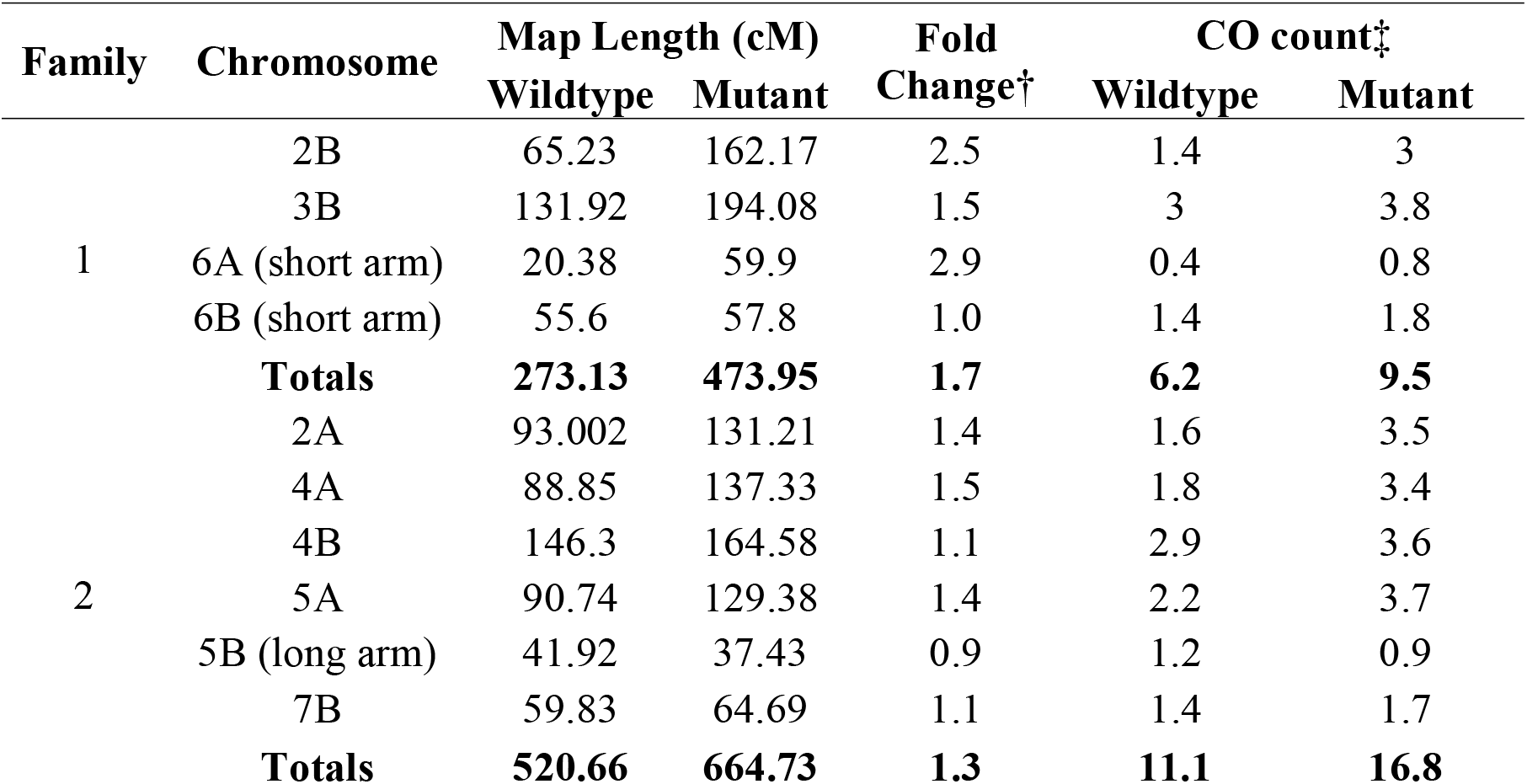
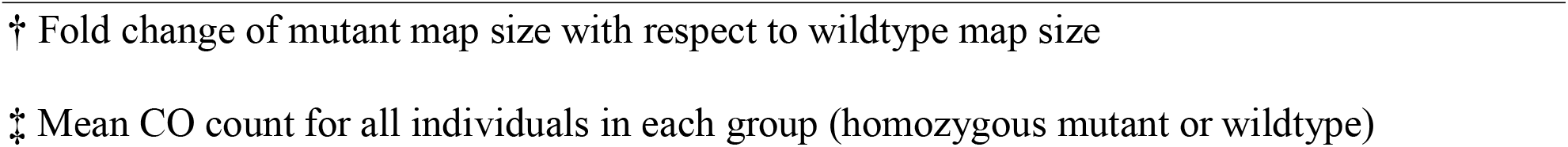
Crossover count and map lengths in lines grouped based on the genotype of the A901T site in *TaTOPII-A1*.

| Family | Chromosome | Map Length (cM) |  | Fold Change<sup>†</sup> | CO count<sup>‡</sup> |  |
| --- | --- | --- | --- | --- | --- | --- |
|  |  | Wildtype | Mutant |  | Wildtype | Mutant |
| 1 | 2B | 65.23 | 162.17 | 2.5 | 1.4 | 3 |
|  | 3B | 131.92 | 194.08 | 1.5 | 3 | 3.8 |
|  | 6A (short arm) | 20.38 | 59.9 | 2.9 | 0.4 | 0.8 |
|  | 6B (short arm) | 55.6 | 57.8 | 1.0 | 1.4 | 1.8 |
|  | **Totals** | **273.13** | **473.95** | **1.7** | **6.2** | **9.5** |
| 2 | 2A | 93.002 | 131.21 | 1.4 | 1.6 | 3.5 |
|  | 4A | 88.85 | 137.33 | 1.5 | 1.8 | 3.4 |
|  | 4B | 146.3 | 164.58 | 1.1 | 2.9 | 3.6 |
|  | 5A | 90.74 | 129.38 | 1.4 | 2.2 | 3.7 |
|  | 5B (long arm) | 41.92 | 37.43 | 0.9 | 1.2 | 0.9 |
|  | 7B | 59.83 | 64.69 | 1.1 | 1.4 | 1.7 |
|  | **Totals** | **520.66** | **664.73** | **1.3** | **11.1** | **16.8** |

† Fold change of mutant map size with respect to wildtype map size
‡ Mean CO count for all individuals in each group (homozygous mutant or wildtype)

In addition to map length comparison, we counted COs using the previously described approach^12^, which is based on counting the number of changes in the parental allele phases across each chromosome for all individuals in a population. For family 1, the average CO count for the wild-type group was 6.2, compared to 9.5 COs for the mutant group. This represents a statistically significant 53% increase and an additive effect size of 1.7 COs (Figure 3b, *p-value* = 0.014 ANOVA). Family 2 also revealed a significant difference between the groups with a mean value of 11.1 COs for wild-type group and 16.8 COs for the mutant group, yielding an additive effect size of 2.9 COs (Figure 3; *p-value* = 0.022 ANOVA). Overall, these effect sizes were larger than those observed from the natural mutation in the NAM population. This could be due to the stronger phenotypic effect of EMS-induced A901T mutation than the naturally occurring P1241L mutation found in the NAM population based on the SIFT prediction scores (0.02 vs 0.05).

## Discussion

Our study shows that both natural and chemically induced alleles of *TaTOPII-A1* affect the frequency of meiotic COs in wheat. The phenotypic effect of EMS mutation in *TaTOPII-A1* was higher than the effect of natural variant segregating in the NAM population, likely due to the amino acid change with more severe predicted functional consequences on protein function, based on SIFT score. The finding that variation in *TOPII* could affect CO frequency is consistent with the results of a previous study in yeast ^10^ and demonstrates that the *TOPII* function in wheat and yeast is conserved. This observation is different from that in *Arabidopsis*^11^, where no detectable differences in CO rate were found between the mutant and wild-type variants of *TOPII*. This could be associated with differences in the meiotic process in *Arabidopsis* and wheat, which have different genome organization, ploidy levels, and more than 100-fold difference in genome size. Similar species-specific effects of mutations in meiotic genes on CO rate have been observed for anti-recombination genes *RecQ4, FANCM* and *FIGL1*^4,7^. For example, mutation in *FANCM* had a significant impact on CO rate in rice and *Arabidopsis*, increasing COs by 2- and 3-fold, respectively^4,17^. However, mutagenesis of *FANCM* in wheat resulted in only modest 31% increase in CO rate^9^. Compared to *fancm* mutants of wheat^9^, chemical mutagenesis of *TaTOPII-A1* gene copy resulted in a more substantial change (51%) in CO rate in the permissive genetic backgrounds of the NAM population founders.

Previously, it was shown that CO rate correlates with SNP density and based on this observation it was proposed that differences in the effects of mutations in meiotic genes on COs could be attributed to species-specific differences in SNP density^4,18^. However, the levels of SNP density in wheat and *Arabidopsis* are similar^19,20^, suggesting that genetic diversity is unlikely factor contributing to this phenomenon. The recently performed megabase-scale analysis of recombination landscape in *Arabidopsis* supports this conclusion and shows that recombination is largely shaped by the genomic distribution of chromatin features rather than by sequence polymorphisms^21^. The low chromatin accessibility regions that are largely devoid of recombination span most of the wheat genome^22–24^, whereas in *Arabidopsis* the inaccessible chromatin cover only small fraction of genome. Thus, it is possible that sensitivity of meiotic genes to chromatin states might influence their impact on CO rate in species with distinct genome organization.

The previous study revealed the complexity of genetic architecture for CO rate variation, which is defined by multiple loci, with the majority having moderate to small additive effects^12^. Our current results indicate that the effects of natural variants of meiotic genes could be also modulated by epistatic interactions with other meiotic QTL. It is likely that the genetic background-specific changes in the direction of meiotic QTL effects observed across the wheat NAM population^12^ are caused by epistatic interactions similar to that demonstrated for *TaTOPII-A1*. Currently, the effect of genetic background on CO rate in meiotic gene mutants across various species is unknown. So far, only *RecQ4* gene mutants consistently increased recombination rate across several plant species with distinct genome organization and sizes, including *Arabidopsis*, rice, tomato and barley^4,7,8^. Therefore, it is possible that interaction between the *TOPII* and genetic background could also contribute to differences in its effect on CO rate between *Arabidopsis* and wheat.

Search for the combinations of meiotic gene mutants capable of increasing CO rate and altering CO distribution in favor of pericentromeric chromosome regions is underway in multiple crop species^4,7,9^. The natural and mutant variants of the *TOPII* gene reported here are a useful addition to the list of recombination promoting genes for manipulating COs in wheat and possibly other crops. Our study also highlights the role of epistasis among meiotic genes in CO rate variation, suggesting that interaction with genetic background will need to be considered while evaluating the phenotypic effects of meiotic gene variants. This improved understanding of the genetic architecture underlying CO rate variation opens up new possibilities for assembling a favorable combination of natural and mutagenized meiotic gene variants to further increase CO rate.

## Materials and Methods

### Natural variation in *TaTOPII-A1*

Originally, QTL affecting total CO and distal CO number was mapped to the region of chromosome 6A containing the conserved wheat homolog of *TOPII* gene (*TaTOPII-A1*)^12^. The whole exome capture and 90K iSelect markers data^13^ obtained for the parental lines of NAM population was mapped to the Chinese Spring genome^25^ and used to identify sequence variation within the *TaTOPII-A1* gene. One-way ANOVA and *t*-tests were used to test an association between CO counts and SNP variation within *TaTOPII-A1*. The IWB34510 marker from the 90K iSelect assay showed strongest association with CO rate in the NAM18 RIL family. Three additional RIL families (NAM19, NAM20, NAM24) segregating at the IWB34510 site, were tested for association using *t*-tests. To compare across families, CO phenotype of the RILs from the four NAM families (NAM18, NAM19, NAM20, and NAM24) were normalized by removing the family effect. Normalized CO (nCO) rates represent the residual value of the individual mean after subtracting the family effect. nCOs were compared across families using one-way ANOVA to assess the effect of the IWB34510 alleles on nCO rates.

### Epistatic interaction analysis

For each unique QTL identified in the NAM population, the marker with the highest LOD score was selected to run against IWB34510 locus in pairwise epistatic analysis. In addition, the markers previously identified using the stepwise regression (SR) or joint composite interval mapping (JCIM) methods^12^ were also included into the analysis of epistasis (Supplementary Table 1). In total, we tested 30 markers using a two-way ANOVA model nCO ∼ IWB34510 + m2 + IWB34510 *m2 + □, where m2 corresponds to the allelic states of one of the 30 QTL markers, nCO to the normalized CO values after removing the family effect, and □ is error. The epistatic effects were considered significant if IWB34510 *m2 interaction term had *p-* value ≤ 0.05.

### EMS mutant of tetraploid wheat

A tetraploid wheat EMS-mutagenized population developed for cv. Kronos^15^, was screened to identify *TaTOPII-A1* mutants. Based on the analysis EMS mutation effects on gene function annotated using exome capture data, we identified TILLING mutant line T4-2306, which carries a non-synonymous mutation A901T in the *TOPII* gene. The M_4_ seeds of line T4-2306 were grown under standard greenhouse conditions to maturity. The DNA extracted from M_4_ plants was genotyped using KASP marker developed by LGC Genomics for the IWB34510 marker to identify lines that were heterozygous for mutation. Two M_4_ plants were selected to develop M_5_ populations, which were genotyped for estimating CO rates. Leaf tissue from M_4_ and M_5_ plants were sampled at the two-leaf stage, and flash frozen at -80**°**C. DNA was extracted using DNeasy 96 (Qiagen) kit. The resulting DNA was quantified using Picogreen (Life Technologies) and normalized to 20 ng/μl for GBS and adjusted to 50 ng/μl for KASP genotyping.

### KASP assays

To develop KASP assays, heterozygous mutation sites across entire genome were extracted for line T4-2306 using exome capture data generated for Kronos mutant population^15^. Genome-specific KASP assays were designed to these mutant sites. Primers were synthesized by IDT, including adding a HEX and FAM sequence to the 5’-end of each wild-type and mutant primer pair, respectively. Parental and progeny DNA were genotyped to identify segregating variants within each family. A total of 112 KASP markers (Supplementary Table 2) were tested across 6 chromosomes for segregation in the two M_5_ families. In total, we detected 15 functional KASP markers segregating in family 1 and 21 functional KASP markers segregating in family 2. The sequences of segregating KASP primers were mapped to the wild emmer reference genome WEW v.2^26^ in order to get a physical position to facilitate genetic map construction. To increase the density of SNP marker data, we used GBS approach for genotyping two M_5_ families.

### Sequence-based genotyping using the GBS approach

GBS was performed on two groups of lines that were selected from the M_5_ generation families to be homozygous for the wild-type and mutant alleles at the A901T site in the *TaTOPII-A1* gene. DNA from each of the 51 selected lines were barcoded and multiplexed in a single GBS library created using the *PstI* and *MseI* restriction enzymes^27^. To reduce library complexity, the genomic library was size-selected with Pippin Prep (Sage Scientific) in the 1.5% gel to retain fragments approximately 300 bp-long, implementing tight size range (261-339 bp).

Two separate runs on Illumina NextSeq instrument were performed at Kansas State University Integrated Genomics Facility, generating 881,071,133 reads. Raw sequencing reads were processed using Illumina fastQC pipeline, resulting in 846,934,516 high quality reads. Quality processed reads were barcode separated using in-house GBS pipeline^27^, resulting in nearly 704 million reads and producing nearly 13.8 million reads per individual line. The reads were aligned to the wild emmer reference genome WEW v.2^26^. The alignments were processed using the Samtools and Picard tools to retain only uniquely mapped alignments. The GATK unified genotyper^28^ was used to call variant sites in the alignments. The raw variant calls were filtered using a conservative criteria to produce high-quality genotype calls for the M_5_ generation lines: 1) only those sites were retained that were homozygous in Kronos DNA sample (wild-type) and heterozygous in the two M_4_ mutant parents DNA sample; 2) the wild-type and mutant alleles in the M_5_ population should follow 1:2:1 segregation ratio; 3) a read depth coverage was filtered to have ≥4 reads; 4) only canonical EMS mutation sites (C->T, G->A) were retained; 5) retained sites must have a genotype calls present in ≥80% of M_5_ generation lines. This stringent filtering retained 50 sites in family 1, and 109 sites in family 2.

### Genetic map construction

The genotyping data generated using the GBS pipeline and KASP assays were combined and input into the ICIM QTL mapping software to construct linkage maps^29^. Due to all informative markers being in the heterozygous state in M_4_ generation plants, the M_5_ progeny genotyping data is expected to segregate as in F_2_ population. Therefore, population type was specified as F_2_ in the ICIM software. Linkage groups and maps were generated using anchor information that was extracted from the physical position of marker in the wild emmer reference genome^26^. Generated maps were rippled using the sum of adjacent distances and final map sizes and plots are reported in cM units (Supplementary Figures 1 and 2). Comparisons of map lengths were made between the wild-type (*TOPII-A1/TOPII-A1*) progeny and mutant (*topII-A1/topII-A1*) progeny for all chromosomes that have at least 5 markers randomly spaced along chromosome or chromosome arm. This resulted in testing 45 markers on 4 chromosomes in family 1 (2B, 3B, short arms of 6A and 6B), and 103 markers on 6 chromosomes in family 2 (2A, 4A, 4B, 5A, 7B, long arm of 5B).

### CO counting

A custom Perl script was written to count the number of COs based on allele phase change from homozygous to heterozygous along each chromosome for the same chromosomes used in ICIM mapping that generated the genetic maps^12^. Results were grouped and distributions were compared between homozygous wildtype *TOPII-A1* allele and homozygous mutant *topII-A1* allele using *t*-test and ANOVA. Allelic additive effects of the mutation were identified as the difference in observed CO number for mutant compared to the wildtype divided by two, [(Hom MUT – Hom WT)/2], and are also reported as percentage change with respect to the wild-type CO number.

## Acknowledgements

This work was funded by the BARD award IS-5116-18 and by the Agriculture and Food Research Initiative Competitive Grants 2022-68013-36439 (WheatCAP). We would like to thank Jorge Dubcovsky for proving seeds of the Kronos mutant lines. Mention of tradenames or commercial products in this publication is solely for the purpose of providing specific information and does not imply recommendation or endorsement by the U. S. Department of Agriculture. USDA is an equal opportunity provider and employer.

