## Supplementary Information for "Allelic variation in the wheat homolog of topoisomerase *II* is associated with crossover rate"

**Supplementary Figures**

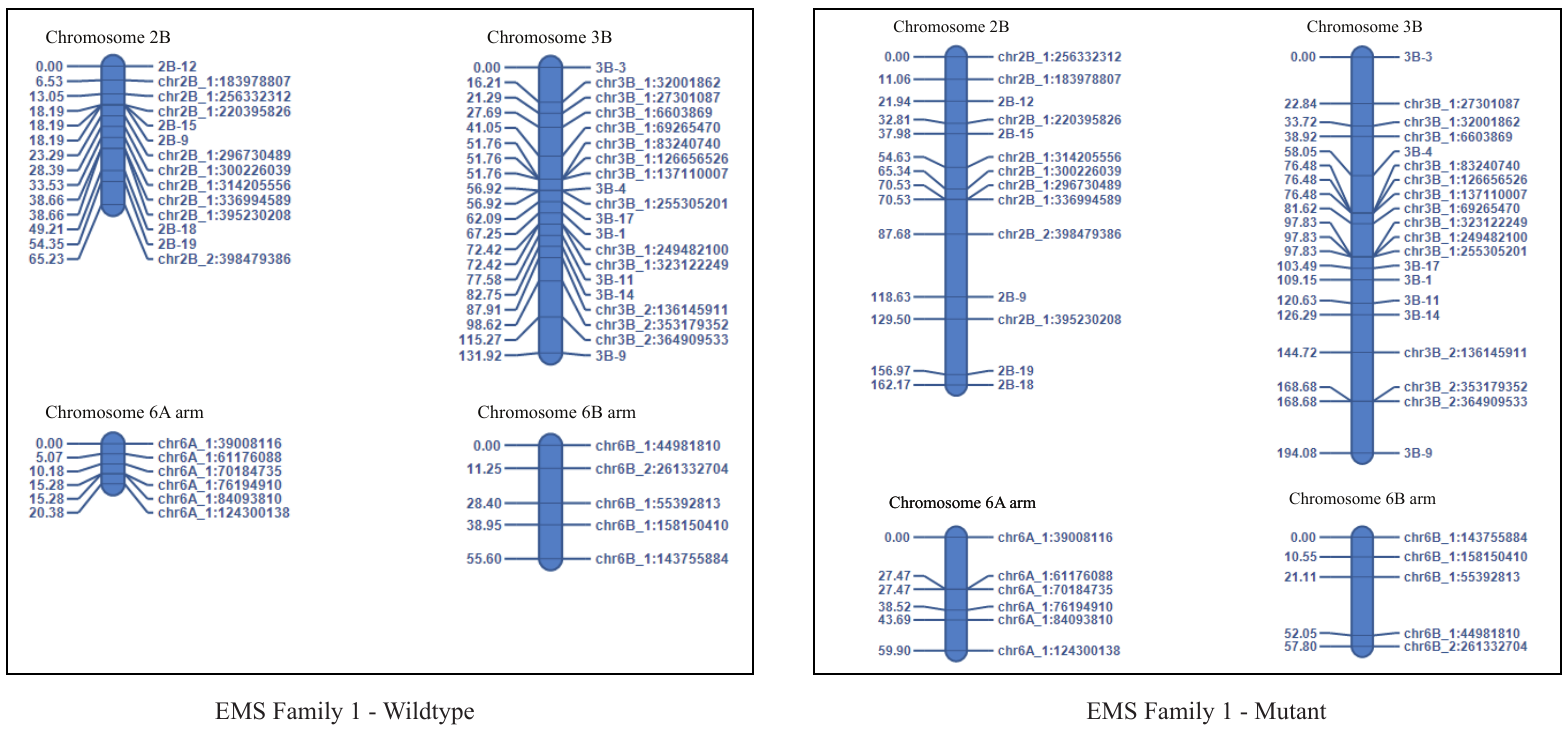

**Supplementary Figure 1.** Comparison of map lengths in family 1 between mutant and wild type alleles of *TaTOPII-A1*

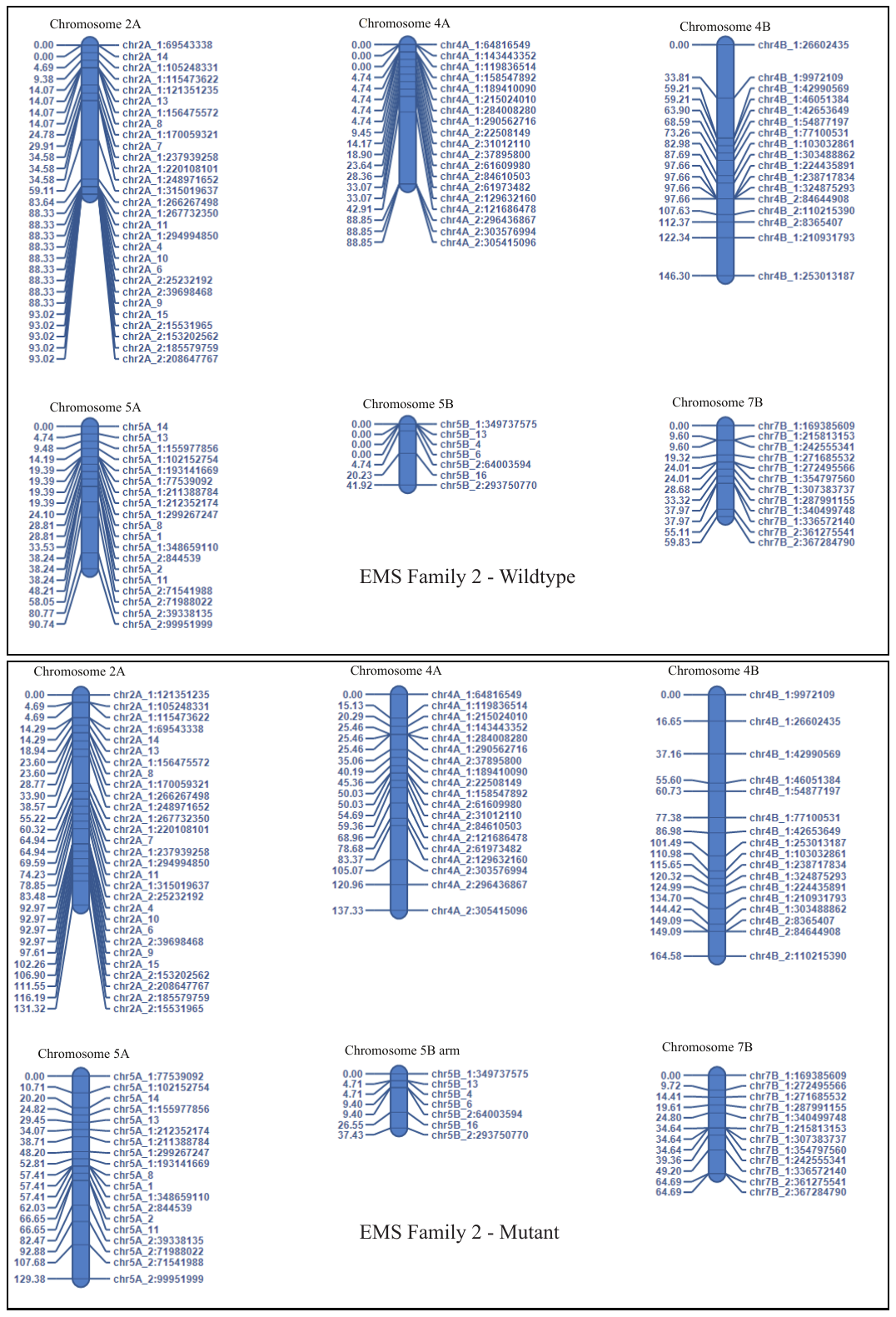

**Supplementary Figure 2.** Comparison of map lengths in family 2 between mutant and wild type alleles of *TaTOPII-A1*

**Supplementary Tables**

| **Supplementary Table 1.** Epistatic interaction analysis between IWB34510 and 30 markers showign association with QTL for CO rate. | | |
| --- | --- | --- |
| **Significant Marker Name (NAM)** | **Significant SNP marker names from 90K SNP array** | **P-value Interaction** |
| BobWhite_c2514_109 |  | 0.42955 |
| BobWhite_c8906_83 |  | 0.37783 |
| BS00022489_51 |  | 0.901 |
| BS00024786_51 |  | 0.2271 |
| BS00105036_51 | IWB12235 | 0.03388 |
| Excalibur_c3423_1170 |  | 0.1488 |
| Excalibur_c57713_81 |  | 0.7826 |
| Excalibur_c72517_227 |  | 0.0775 |
| Excalibur_c84439_196 |  | 0.6652 |
| Kukri_c32253_407 |  | 0.445 |
| Kukri_c33670_261 |  | 0.494 |
| Kukri_c59320_140 |  | 0.239 |
| Kukri_rep_c109167_89 |  | 0.53 |
| Ra_c35412_494 |  | 0.386 |
| RAC875_c716_198 |  | 0.605 |
| RAC875_rep_c105584_237 |  | 0.206 |
| RAC875_rep_c72220_143 |  | 0.0275 |
| RAC875_rep_c82932_407 |  | 0.759 |
| SpringWheatNAM_tag_100953 |  | 0.395 |
| SpringWheatNAM_tag_102286 |  | 0.6655 |
| SpringWheatNAM_tag_121667 |  | 0.1709 |
| SpringWheatNAM_tag_17106 |  | 0.6817 |
| SpringWheatNAM_tag_177026 |  | 0.008551 |
| SpringWheatNAM_tag_236732 |  | 0.783 |
| SpringWheatNAM_tag_256152 |  | 0.001228 |
| SpringWheatNAM_tag_49013 |  | 0.4025 |
| SpringWheatNAM_tag_64311 |  | 0.1877 |
| Tdurum_contig30426_261 |  | 0.9376 |
| tplb0055o21_1994 | IWB75092 | 0.001505 |
| wsnp_Ex_c19773_28772235 |  | 0.303 |

**Supplementary Table 2.** List of markers used for counting COs in the populations derived from the Kronos *TaTOPII-A1* mutant T4-2306.
